## Supplementary figures for "Filamentation of the bacterial bi-functional alcohol/aldehyde dehydrogenase AdhE is essential for substrate channeling and enzymatic regulation"

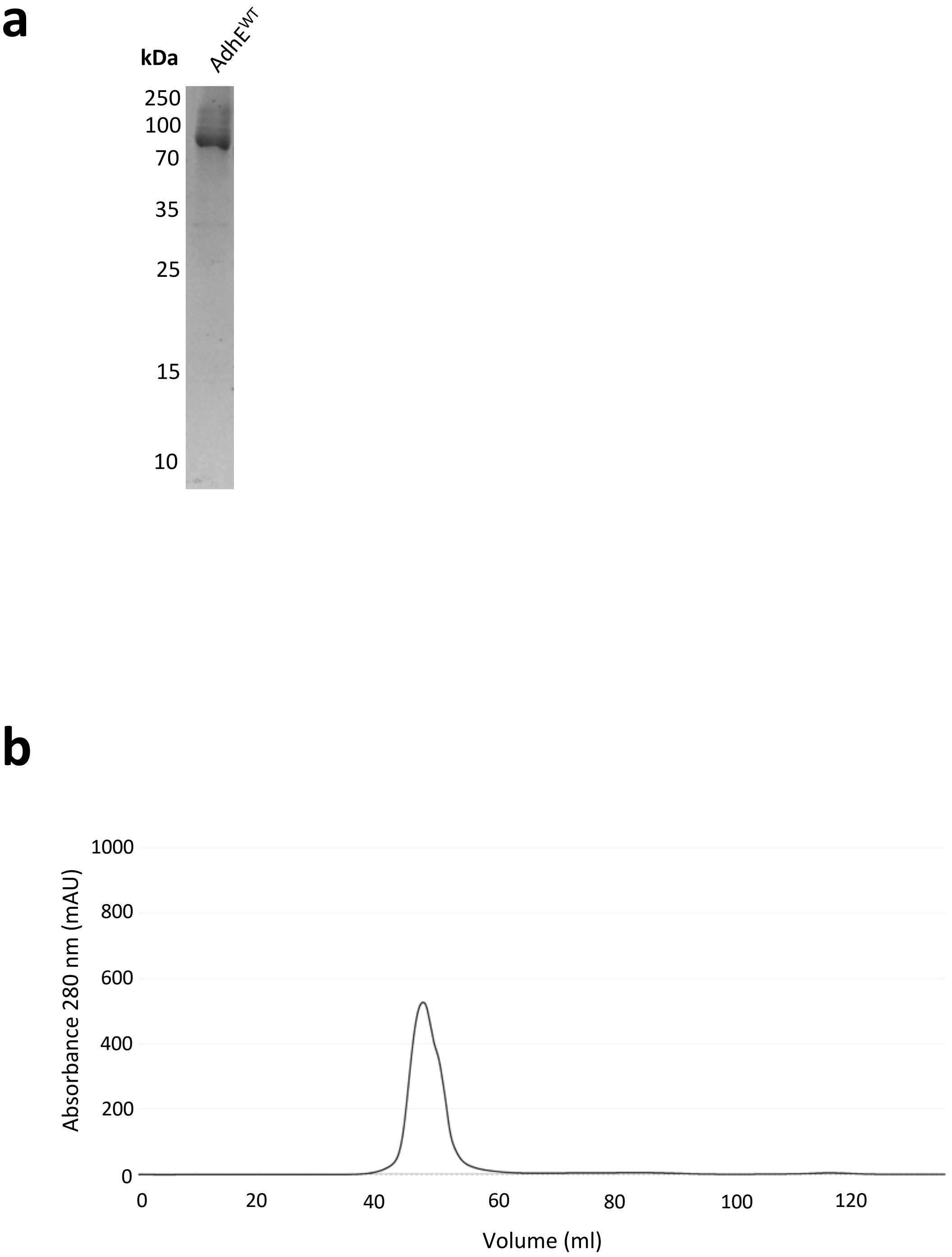


**Supplementary Fig. 1. Purification of spirosomes. a.** SDS-PAGE (12%) of purified AdhE$^{WT.}$ Molecular weight are indicated on the left. **b.** Size-exclusion chromatography profile of purified AdhE$^{WT}$ on a HiLoad 16/600 Superdex 200 pg column.

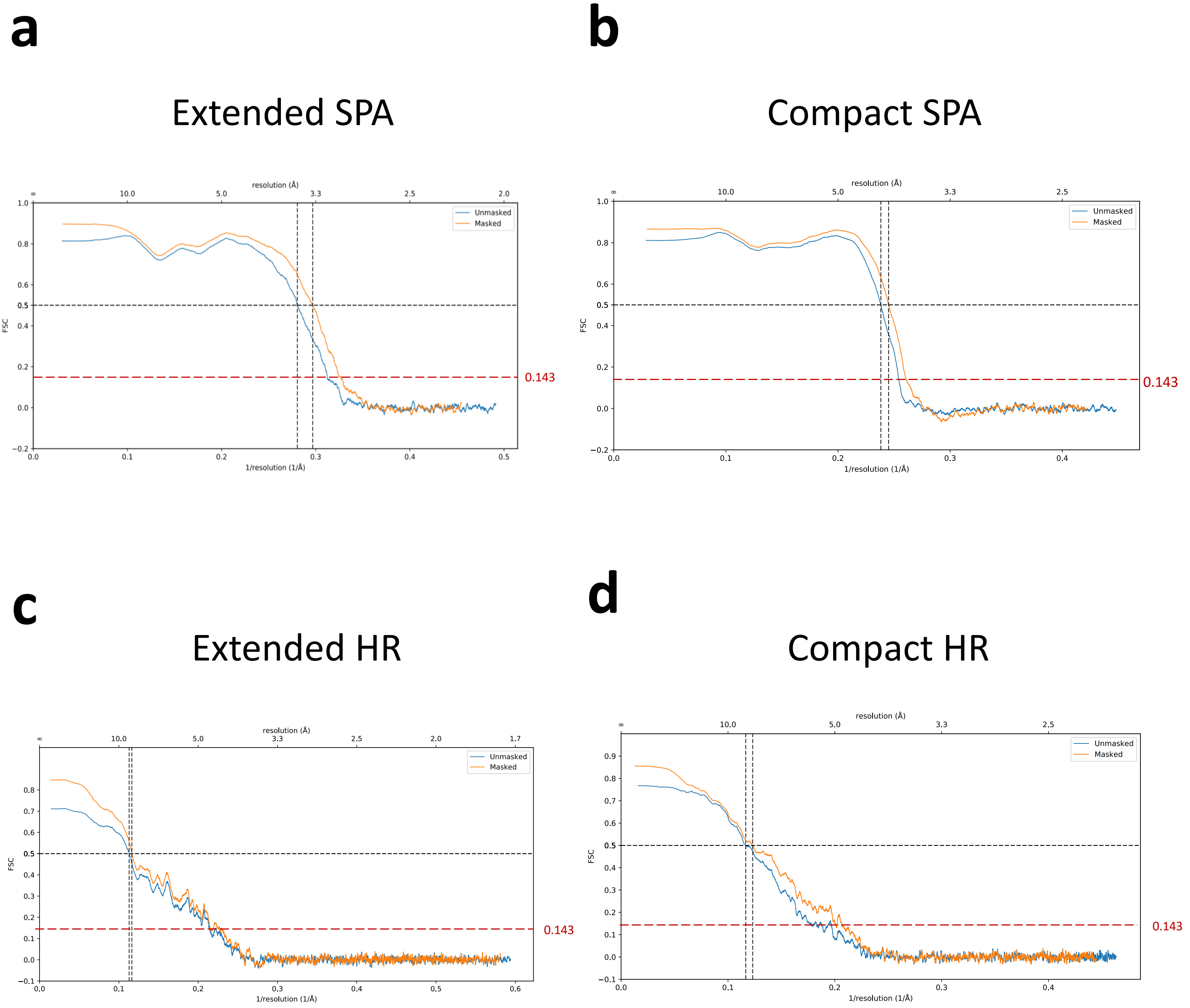


**Supplementary Fig. 2. Resolution of the spirosome cryo-EM maps.** Fourier shell correlation (FSC) curves for masked and unmasked reconstructions. **a.** extended spirosome obtained using SPA **b.** compact spirosome obtained using SPA **c.** extended spirosome obtained using HR **d.** compact spirosome obtained using HR. The resolution was calculated at the cut-off 0.143 of the FSC (red dashed line).

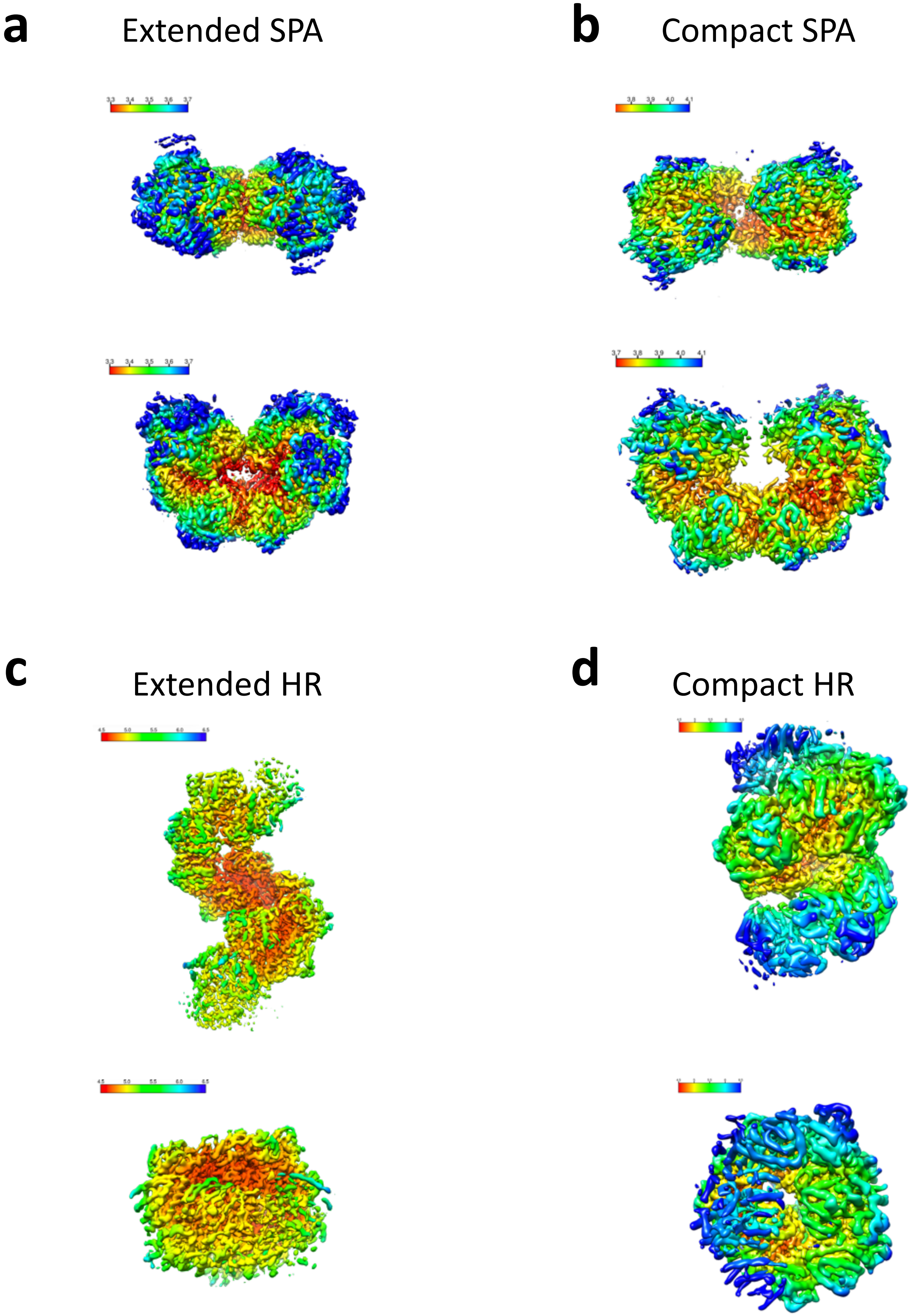


**Supplementary Fig. 3. Representation of the local resolution in the cryo-EM maps .** Local resolution is displayed on the cryoEM maps. A color-coded scale indicates the resolution values dispayed for each map **a.** extended spirosome (3.3Å (red) to 3.7Å (blue) resolution) obtained using SPA **b.** compact spirosome (3.7Å (red) to 4.1Å (blue) resolution) obtained using SPA. **c.** and **d.** extended spirosome and compact spirosome (4.5Å (red) to 6.5Å (blue) resolution) obtained using HR.

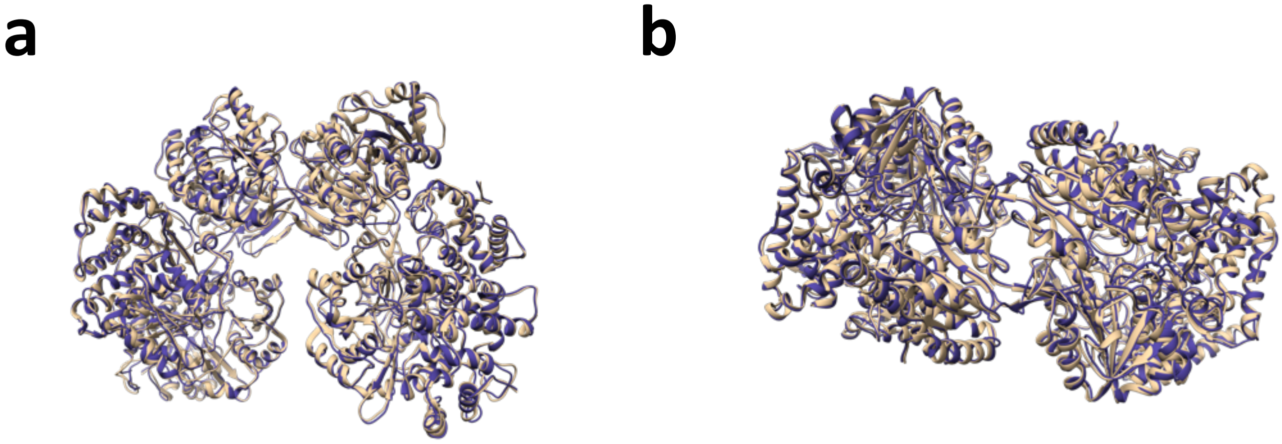


**Supplementary Fig. 4. Comparison of apo and compact spirosome.** Superimposition of the structures of the apo (PDB code: 6AHC) and the spirosomes in presence of NADH and $Fe^{2+}$.

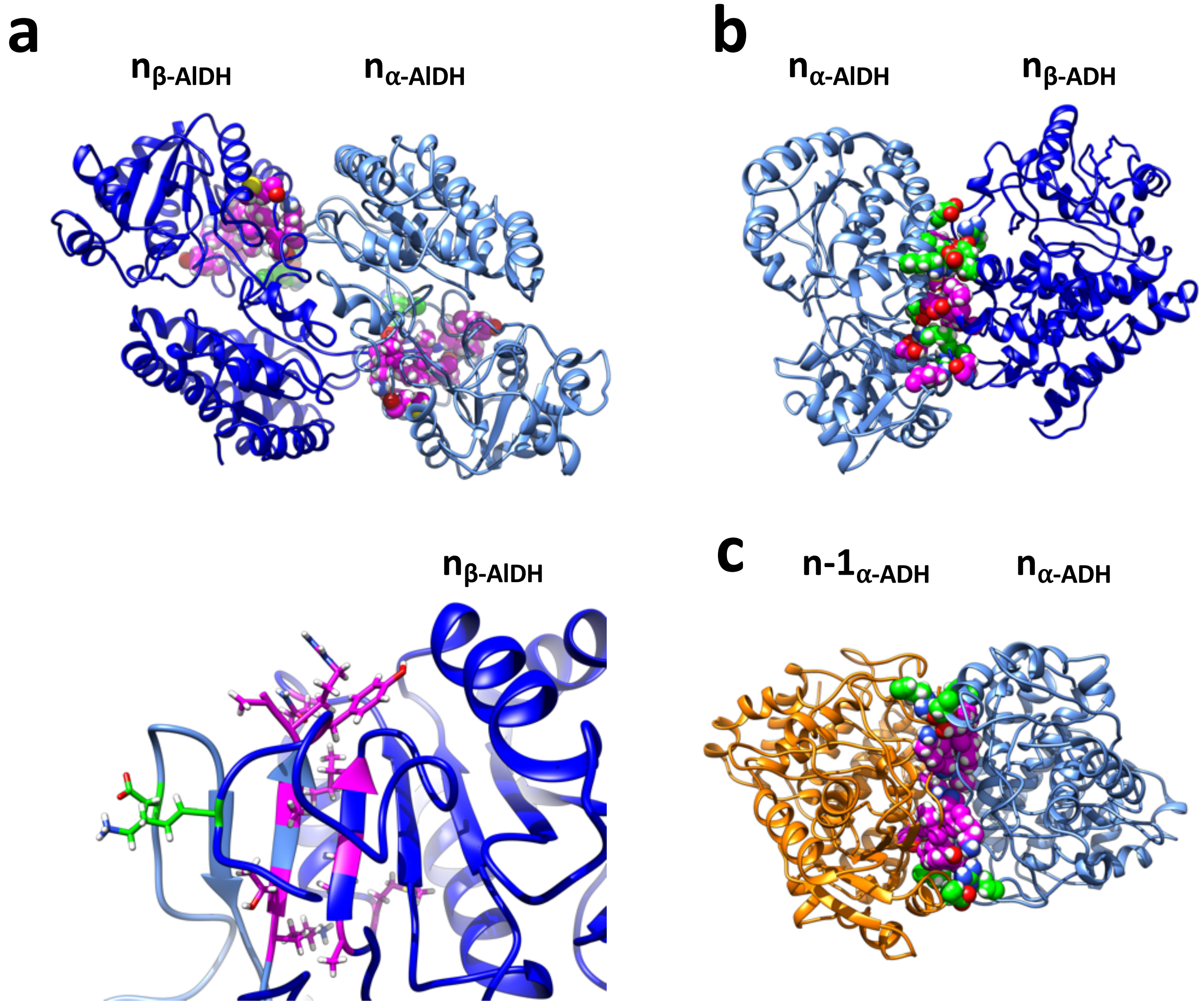


**Supplementary Fig. 5. Representation of the residues involved in inter-domain interfaces within the extended spirosome.** **a**. Ribbon representation of the acetaldehyde $n_\beta$ (dark blue) – acetaldehyde $n_\alpha$ (light blue) dehydrogenase. Lower panel, zoomed representation of the main interface (beta-complementation) between the acetaldehyde $n_\beta$ (dark blue) – acetaldehyde $n_\alpha$ (light blue) dehydrogenase **b.** Ribbon representation of the acetaldehyde $n_\alpha$ (light blue) – alcohol $n_\beta$ (dark blue) dehydrogenase **c.** Ribbon representation of the alcohol $n\text{-}1_\alpha$ (orange) – alcohol $n_\alpha$ (light blue) dehydrogenase interfaces.

Residues involved in hydrogen bonds are displayed as spheres in pink and residues involved in salt bridges are displayed as spheres in green.

**a**

| | ALDH | ADH | BOND |
|---|---|---|---|
| AlDH – ADH interface | GLU 63 | TYR 848 | H |
| | | ILE 851 | H |
| | HIS 70 | ARG 572 | H |
| | GLU 74 | MET 574 | H |
| | ARG 229 | SER 764 | H |
| | THR 239 | LYS 759 | H |
| | LYS 317 | PRO 757 | H |
| | ASP 44 | ARG 572 | SB |
| | ARG 46 | ASP 838 | SB |
| | LYS 51 | GLU 834 | SB |
| | ASP 64 | LYS 759 | SB |
| | ARG 229 | ASP 767 | SB |

**b**

| | AlDH | AlDH | BOND |
|---|---|---|---|
| AlDH – AlDH interface | GLN 375 | ARG 447 | H |
| | GLY 386 | LYS 442 | H |
| | MET 389 | | H |
| | ALA 392 | THR 443 | H |
| | ILE 394 | ALA 445 | H |
| | ILE 396 | ARG 447 | H |
| | TYR 409 | ALA 448 | H |
| | SER 430 | LYS 441 | H |
| | ASP 92 | LYS 412 | SB |

**c**

| | ADH | ADH | BOND |
|---|---|---|---|
| ADH – ADH interface | LEU 452 | PHE 462 | H |
| | HIS 454 | ILE 460 | H |
| | LYS 455 | TYR 461 | H |
| | GLU 669 | SER 705 | H |
| | PHE 670 | GLN 674 | H |
| | LYS 455 | GLU 473 | SB |
| | ARG 463 | GLU 669 | SB |
| | GLU 473 | LYS 455 | SB |
| | | ARG 577 | SB |

**Supplementary Fig. 6. PISA analysis of the 3 interfaces.** Residues involved in hydrogen bonds and salt bridges are marked H and SB respectively. **a.** acetaldehyde (AlDH) – alcohol (ADH) dehydrogenase interface, **b.** acetaldehyde (AlDH) – acetaldehyde (AlDH) dehydrogenase interface **c.** alcohol (ADH) – alcohol (ADH) dehydrogenase interface.

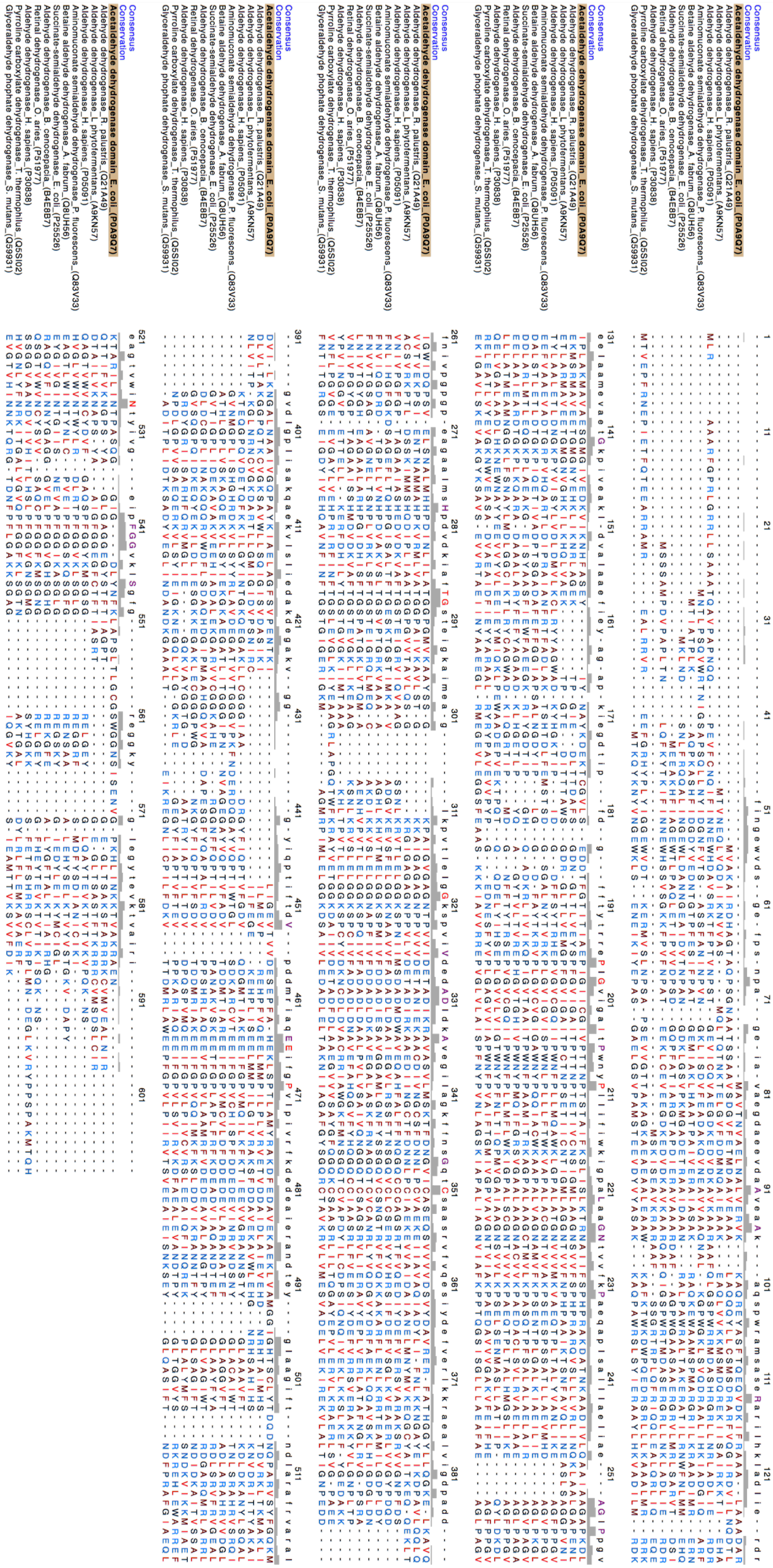

**Supplementary Fig. 7. Multiple sequence alignement of AlDH domain of *E. coli* AdhE and monofunctional AlDH enzymes.**

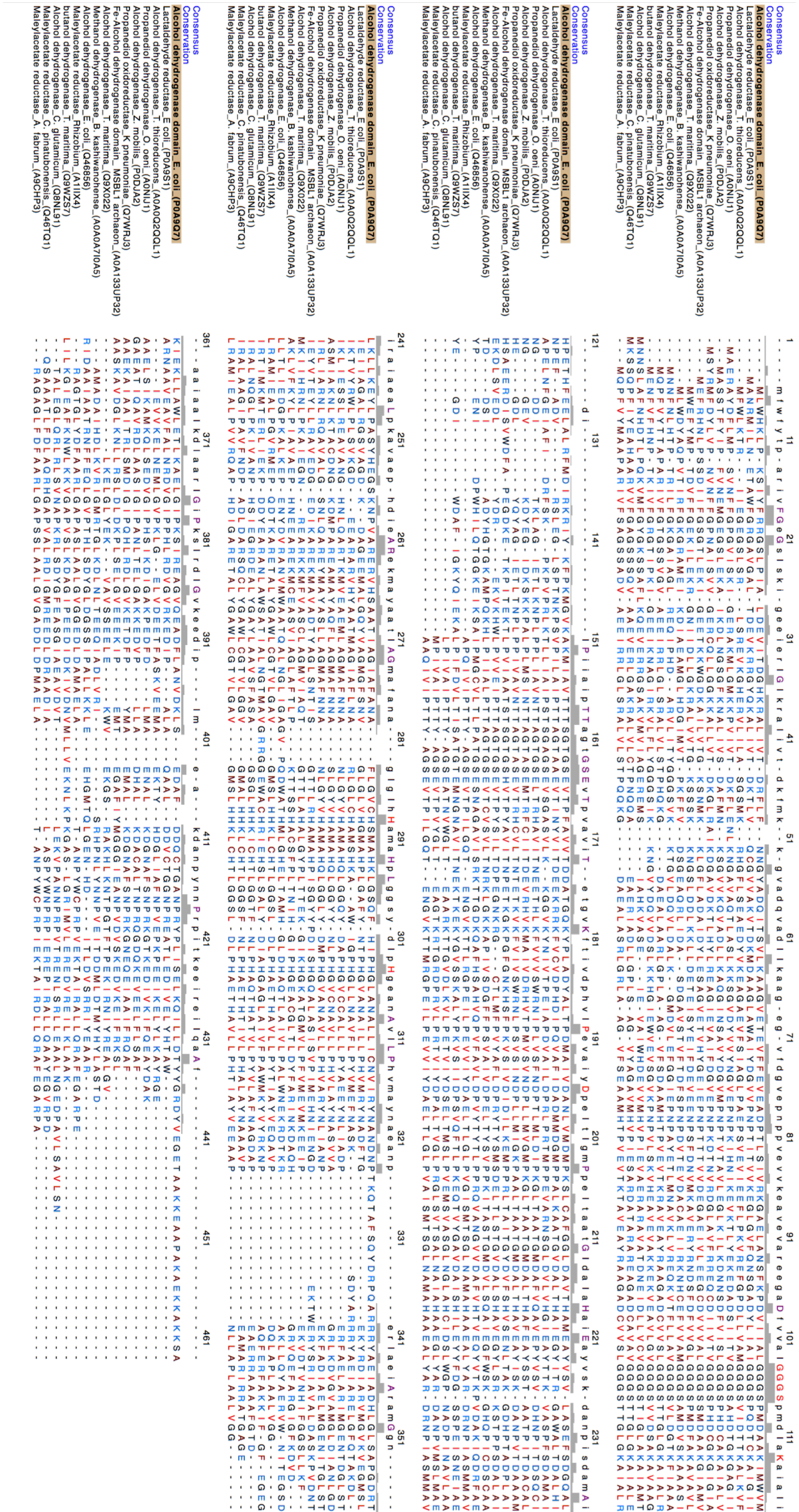

**Supplementary Fig. 8. Multiple sequence alignement of ADH domain of *E. coli* AdhE and monofunctional ADH enzymes.**

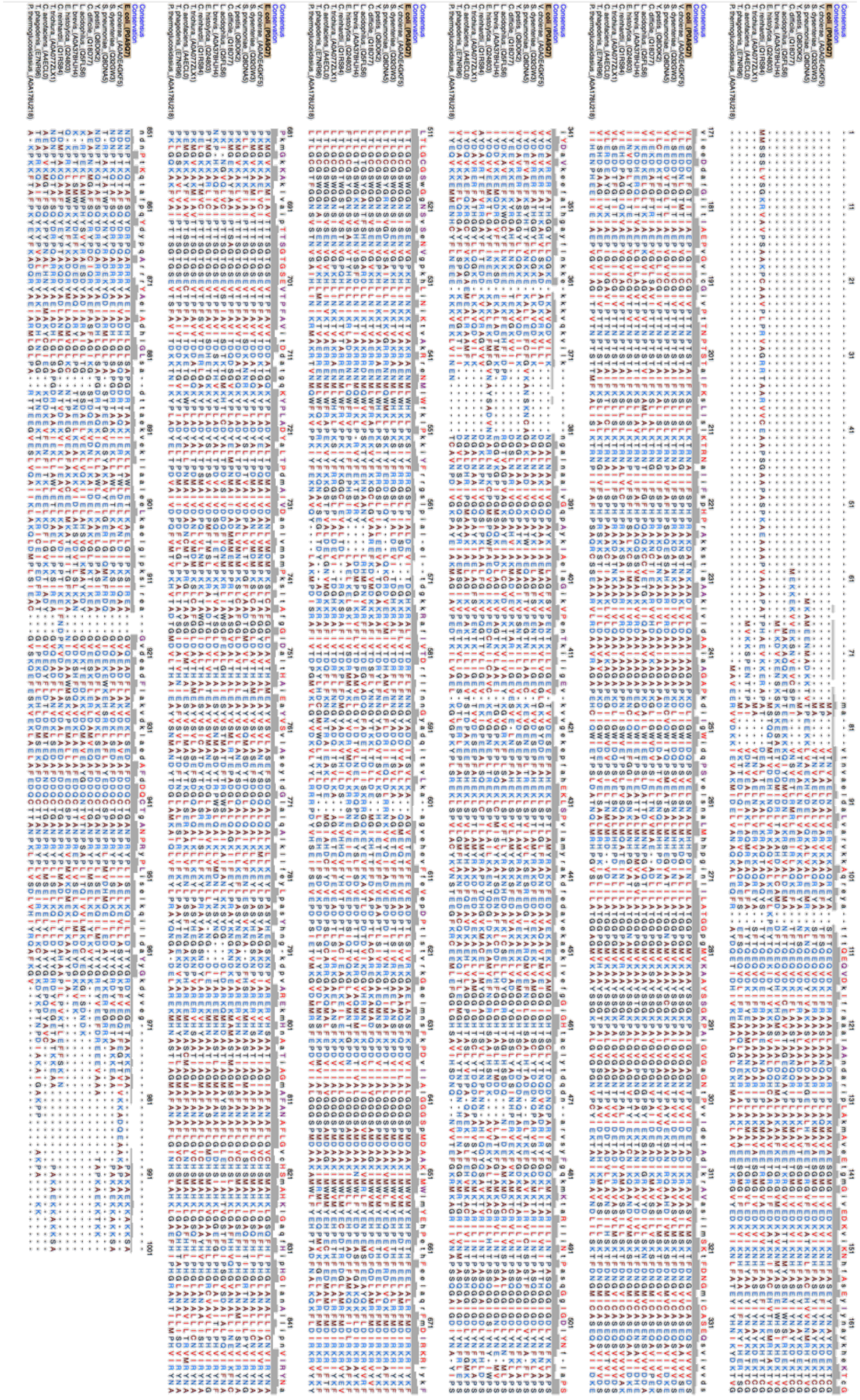

**Supplementary Fig. 9. Multiple sequence alignement of *E. coli* AdhE and oher AdhE homologs.**

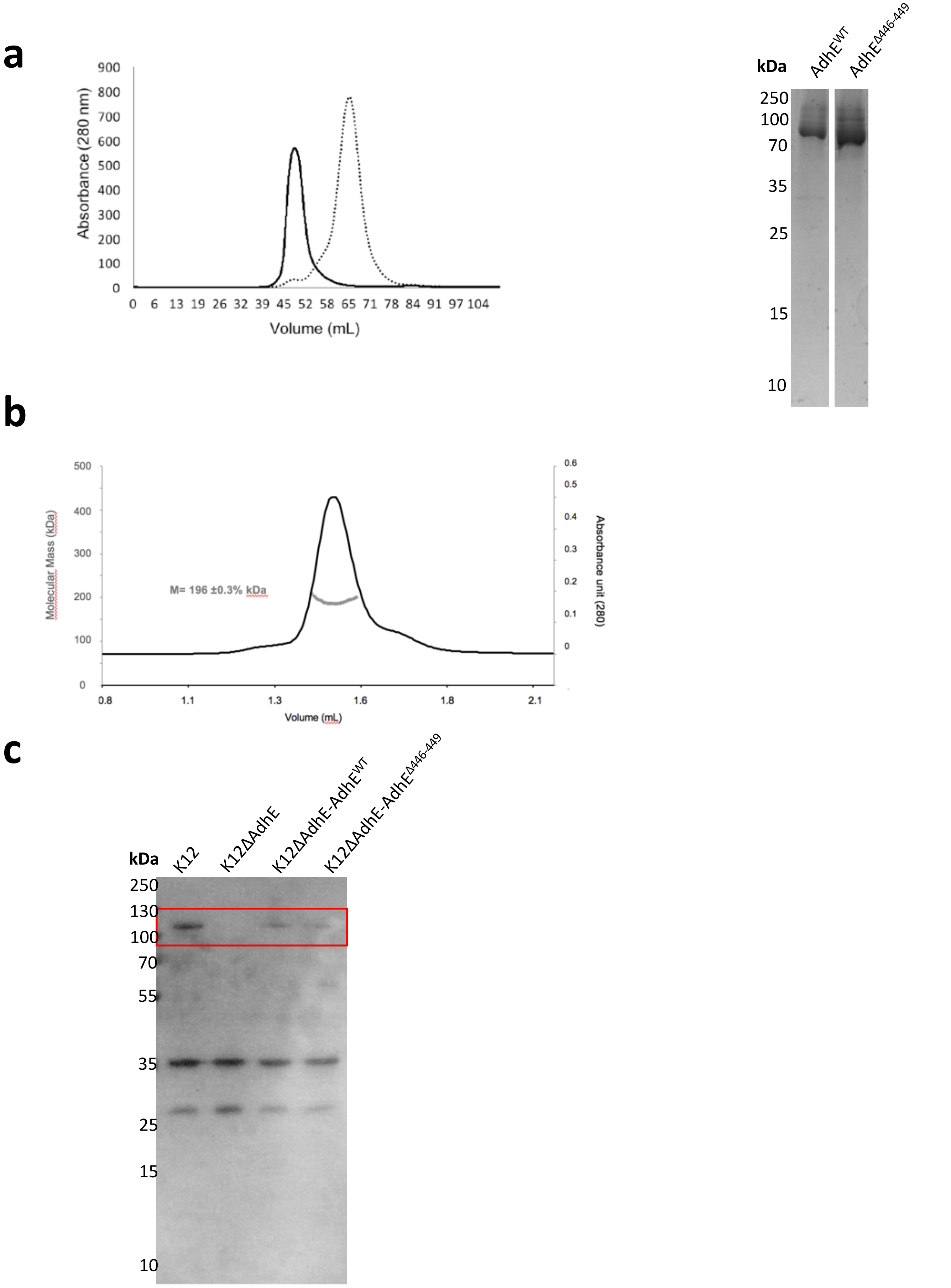


**Supplementary Fig. 10. Expression test and purification of AdhE$^{WT}$ and AdhE$^{\Delta 446\text{-}449}$. a.** Size-exclusion chromatography profile of purified AdhE$^{WT}$ (dark line) and AdhE$^{\Delta 446\text{-}449}$ (dashed line) on a HiLoad 16/600 Superdex 200 column (left) and SDS-PAGE (12%) of AdhE$^{WT}$ and AdhE$^{\Delta 446\text{-}449}$ (right). Molecular weight markers are indicated on the left. **b.** Size-exclusion profile of purified AdhE$^{\Delta 446\text{-}449}$ on a Superdex S200 5/150 column coupled with MALS and RI. **c.** Western-blot using anti-AdhE antibody. Molecular weight markers are indicated on the left.
